# Crystal structure of the kinase domain of a receptor tyrosine kinase from a choanoflagellate, *Monosiga brevicollis*

**DOI:** 10.1101/2022.10.07.511304

**Authors:** Teena Bajaj, John Kuriyan, Christine L. Gee

## Abstract

Genomic analysis of the unicellular choanoflagellate, *Monosiga brevicollis* (MB), revealed the remarkable presence of cell signaling and adhesion protein domains that are characteristically associated with metazoans. Strikingly, receptor tyrosine kinases, one of the most critical elements of signal transduction and communication in metazoans, are present in choanoflagellates. We determined the crystal structure at 1.95 Å resolution of the kinase domain of the *M. brevicollis* receptor tyrosine kinase C8 (RTKC8, a member of the choanoflagellate receptor tyrosine kinase C family) bound to the kinase inhibitor staurospaurine. The chonanoflagellate kinase domain is closely related in sequence to mammalian tyrosine kinases (∼ 40% sequence identity to the human Ephrin kinase domain EphA3) and, as expected, has the canonical protein kinase fold. The kinase is structurally most similar to human Ephrin (EphA5), even though the extracellular sensor domain is completely different from that of Ephrin. The RTKC8 kinase domain is in an active conformation, with two staurosporine molecules bound to the kinase, one at the active site and another at the peptide-substrate binding site. We show that the RTKC8 kinase domain can phosphorylate tyrosine residues in peptides from its C-terminal tail segment.

## Introduction

Receptor tyrosine kinases are crucial signaling molecules in animal cells, regulating inter-cellular responses in various ways. These receptors consist of an extracellular ligand-binding domain, a single transmembrane helix, and a cytoplasmic tyrosine kinase domain [1,2]. The binding of extracellular ligands to the receptor leads to activation of the kinase domain, which then autophosphorylates tyrosine residues present in a centrally located activation loop in the kinase domain, in a C-terminal tail of the receptor, or in loops in the kinase domain, or in the juxtamembrane segment bridging the transmembrane helix and the kinase domain. These phosphorylated tyrosine residues switch on kinase activity, or act as docking sites for proteins that transmit the signal further downstream [3–5].

The discovery of numerous genes encoding receptor tyrosine kinases in choanoflagellate genomes, was remarkable [6–10]. Phylogenetic studies suggest that choanoflagellates are the closest living relatives to metazoans, indicating that receptor tyrosine kinases were present in the common ancestor of choanoflagellates and metazoans. The choanoflagellate *M. brevicollis* contains 128 protein kinases, including 88 receptor tyrosine kinases [11]. The choanoflagellate receptor tyrosine kinases are classified into 15 families (denoted A through M, FGTK and LRTK) [11]. These tyrosine kinases are not direct orthologs of metazoan receptor tyrosine kinases as they have extracellular domains that appear unrelated to those of metazoan receptor tyrosine kinases [11]. The structural and functional aspects of choanoflagellate receptor tyrosine kinases are still relatively unexplored.

The *M. brevicollis* receptor tyrosine kinase C family has 10 members (denoted RTKC1 to RTKC10). The RTKC family is a distinct clade of kinases and shares a common ancestor with the Ephrin kinases but does not share all of the Ephrin-specific conserved motifs [12]. Most of the members of this family contain Cys-rich and Hyalin-rich (HYR) extracellular domains, a transmembrane domain, a kinase domain and a C-terminal tail with a CAP-Gly domain. The HYR domain is predicted to be related to immunoglobulin (Ig) and fibronectin type3 (FN3) domains in metazoans, and might be involved in cell adhesion [11,13]. The metazoan CAP-Gly domains bind to microtubules [14]. The presence of a CAP-Gly domain in *M. brevicollis* C family receptor tyrosine kinases therefore suggests that these kinases may interact with the cytoskeleton [11]. *M. brevicollis* RTKC8 has an extracellular domain of ∼1500 residues containing two Cys-rich domains and four HYR domains. The intracellular domain consists of a kinase domain (∼280 residues) and a C-terminal tail containing a 40 residue CAP-Gly domain and 10 tyrosine residues (Fig 1A).

**Fig 1.**
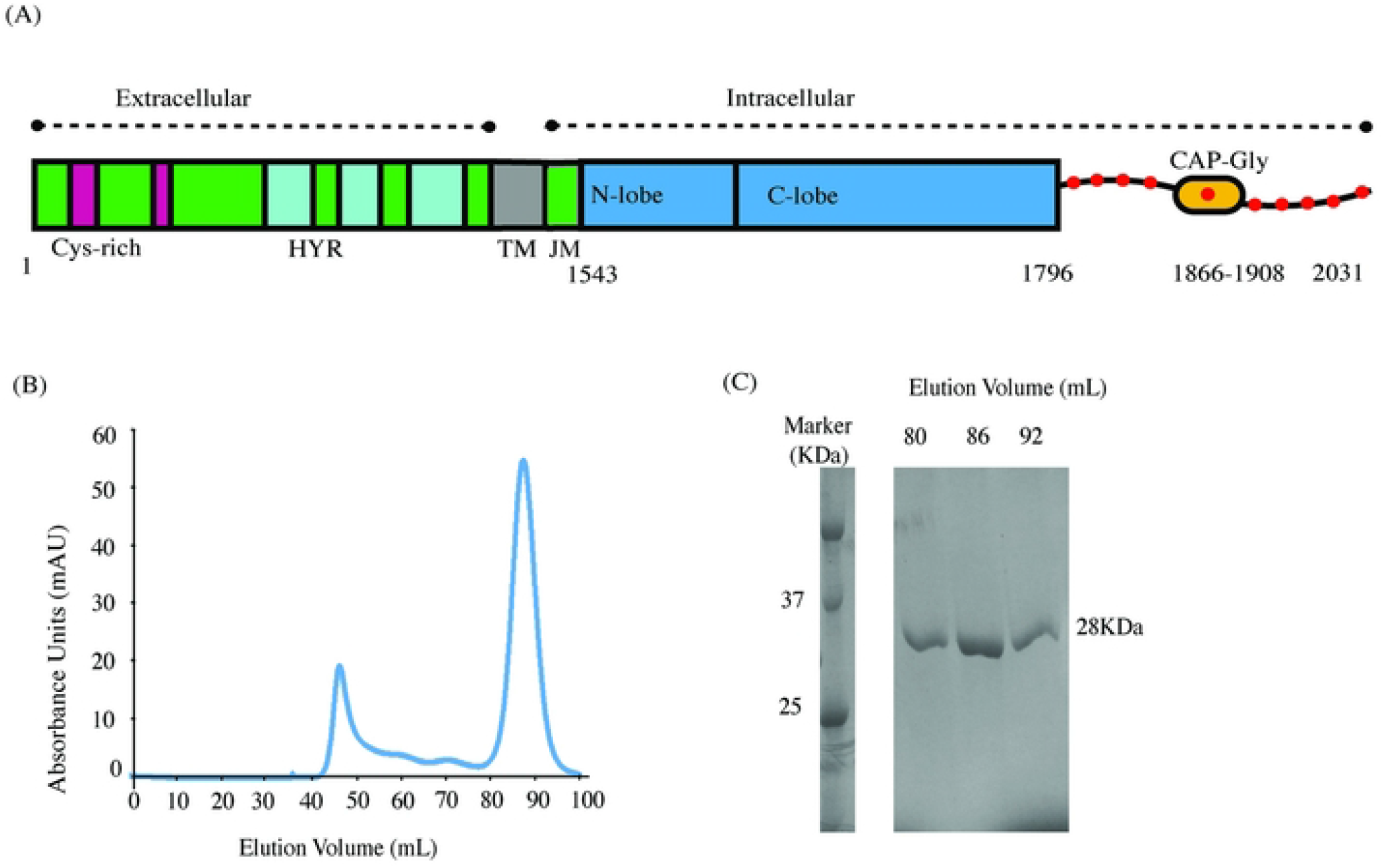
Architecture of the RTKC8 receptor tyrosine kinase and purification of its kinase domain. (A) Domain organization of full length RTKC8, consisting of Cys-rich and hyalin rich (HYR) domains in the extracellular domain, followed by a transmembrane domain and an intracellular domain. The intracellular domain contains a juxtamembrane domain, the kinase domain, the C-terminal tail with tyrosine residues (red circles) and a CAP-Gly domain. The expressed construct consists of an N-terminal hexa-His tag followed by a TEV protease site and the cytoplasmic kinase domain (residues 1530-1811). (B) Chromatogram showing the protein eluted from gel filtration as a single symmetric peak at the expected elution volume for a monomer. (C) SDS-gel of fractions from gel filtration confirmed the protein purity.

We determined the crystal structure of the RTKC8 kinase domain to 1.95 Å resolution. The RTKC8 kinase domain is co-crystallized with the kinase inhibitor staurosporine.

## Materials and Methods

### Cloning of the RTKC8 kinase domain

The sequence of the RTKC8 kinase matches that reported in the Kinase.com database (http://kinase.com/kinbase) [11]. The DNA for the kinase domain without the transmembrane or tail segment (residues 1530-1811) of RTKC8 was amplified using standard PCR protocols from a Monosiga cDNA library (a gift from Nicole King) using a forward primer (5’AGGGCCATATGCAGCTTTCCAAGGAGCCGCGCG3’) and a reverse primer (5’CCTTTGAATTCTCAGCTTTGGAAGAGGCTGGACATGTCCGAATCAGT3’). The PCR product size was confirmed using agarose gel electrophoresis. The amplified DNA was ligated into the pFastbac1 insect-cell expression vector using EcoRI (5’) and NdeI (3’) restriction enzyme sites. The ligated product was transformed into *E. coli* Top10 chemically competent cells. Next day, the overnight culture was set up from Top10 bacterial colonies to isolate the plasmid and the plasmid was sequenced to confirm the cloned construct. The expressed construct consisted of the *M. brevicollis* receptor tyrosine kinase C8 kinase domain with an N-terminal hexahistidine tag and a TEV protease site.

### Expression of the RTKC8 kinase domain

The pFastbac1-RTKC8-kinase vector carrying the ampicillin resistance gene was transformed into DH10bac *E. coli* cells for transposition into the bacmid. The outgrowth (diluted 1:100) was plated over Luria-Bertani (LB) agar plates containing kanamycin (50 µg/mL), gentamycin (7 µg/mL), tetracycline (10 µg/mL), isopropyl β-D-1-thiogalactopyranoside (IPTG) (40 µg/mL), bluo-gal (100-300 µg/mL) and incubated at 37°C. After 24-48 hours, the white colonies (containing the cloned construct) were cultured overnight to isolate the bacmid. The bacmid was transfected into *Spodoptera frugiperda* (Sf9) cells to prepare baculovirus stock, which was further amplified via three cycles of infection. After 3 cycles of virus amplification, the supernatant of this third passage (P3 virus) was used to infect another 2 liter batch of Sf9 cells grown to 2.5 × 106 cells/ mL in the presence of 10% antibiotics (penicillin and streptomycin). Cells were harvested after 48 hours. Each liter of culture was resuspended in 20 mL of lysis buffer (50 mM Tris-HCl pH 8.0, 200 mM NaCl, 20% glycerol, 10 mM CaCl_2_, 10 mM MgCl_2_, 50 mM imidazole, 0.5 mM TCEP) containing 10 µg/mL DNaseI and a protease inhibitor cocktail of 4-(2-Aminoethyl) benzenesulfonyl fluoride hydrochloride (AEBSF) (0.2 mM), benzamidine (0.5 mM) and leupeptin (0.005 mM). The resuspended lysate was stored at -80°C until thawed for purification.

### Purification of the RTKC8 kinase domain

To purify the protein, cells were thawed and lysed using an Avestin EmulsiFlex C50 cell homogenizer. The suspension was clarified by ultracentrifugation at 40,000 rpm for 1 hour at 4°C. The supernatant was filtered using a 0.45 µM syringe filter, loaded onto a Ni-NTA affinity column (GE Healthcare Life Sciences) pre-equilibrated with Buffer A (50 mM Tris-HCl pH 8.0, 500 mM NaCl, 10% glycerol, 25 mM imidazole, 0.5 mM TCEP) and the flow-through was collected. The resin was washed with 20 column volumes of Buffer A to remove the non-specific proteins. Then, the RTKC8 protein was eluted in Buffer B (50 mM Tris-HCl pH 8.0, 200 mM NaCl, 10% glycerol, 250 mM imidazole, 0.5 mM TCEP). To cleave the His-tag, TEV protease was added to the pooled fractions containing the protein and dialyzed into Buffer C (50 mM Tris-HCl pH 8.0, 200 mM NaCl, 10% glycerol, 0.5 mM TCEP) overnight at 4°C. A subtractive Ni-NTA purification was run to remove TEV protease and other impurities. The flow through was collected, concentrated and purified on a Superdex 200 16/600 column (GE Healthcare Life Sciences) in Buffer C. Fractions containing RTKC8 kinase were either pooled and concentrated to 105 µM for biochemical experiments and stored at -80°C or concentrated to 200 µM and used immediately for crystallization experiments. The purification yield was ∼1 mg of purified RTKC8 kinase from 1 liter of Sf9 insect cells. The elution volume from size exclusion was consistent with the molecular weight of a monomeric protein (28.5 KDa) (Fig1B) and ran as a single chromatography band on SDS PAGE (Fig 1C).

### Phosphorylation and dephosphorylation analysis of RTKC8 kinase domain

To determine the phosphorylation state of the purified RTKC8 kinase domain, the protein was probed using an anti-phosphotyrosine antibody (4G10) by western blot. The protein was run on 12% acrylamide SDS-PAGE gel at a final concentration of 1 µM and transferred to polyvinylidene difluoride (PVDF) membrane (Millipore) using a semi-dry transfer apparatus. Protein samples were transferred using Towbin transfer buffer (25 mM tris, 192 mM glycine, 20% methanol). The membrane was blocked using blocking solution (5% BSA dissolved in tris-buffered saline (20 mM Tris pH 7.5, 150 mM NaCl), 0.1% TritonX-100). The membrane was probed using anti-phosphotyrosine 4G10 primary antibody (Millipore, 05-321) (1:3000), diluted in blocking solution overnight at 4°C. Next day, the blot was washed with 1X tris-buffered saline containing 0.1% TritonX-100 for four times for 5 minutes each with gentle shaking at room temperature. The blot was probed using anti-Mouse IgG secondary antibody linked with horse radish peroxidase (Cell Signaling Technology, #7076) (1:3000) for 1 hour at room temperature on shaker. The blot was washed again with 1X Tris Bis solution containing 0.1% TritonX-100 for four times for 5 minutes with gentle shaking at room temperature. The blot was developed using the Western Bright kit (Advansta) and imaged using a Bio-Rad Gel Doc.

For the dephosphorylation assay, the RTKC8 kinase domain and YopH phosphatase, each at a final concentration of 1 µM, were incubated together at room temperature and the dephosphorylation reaction was stopped at different time intervals by adding SDS-buffer. Phosphotyrosine levels were detected by western blot as described above.

### *In vitro* kinase activity assays

Kinase activity measurements were carried out using a coupled kinase assay (Pyruvate Kinase/Lactate Dehydrogenase (PK/LDH) ATPase assay) in presence of different substrate peptides [15,16]. This assay measures ADP production, which is coupled to NADH oxidation. In this experiment, the reaction solution contained 50 mM Tris-HCl pH 8, 100 mM NaCl, 10 mM MgCl_2_, 1 mM phosphoenolpyruvate (PEP), 300 mg/mL NADH, 2 mM sodium orthovanadate (Na_3_VO_4_), 100 µM ATP, and an excess of pyruvate kinase and lactate dehydrogenase (approximately 120 units/mL and 80 units/mL, respectively, added from a commercially available mixture of the two enzymes). Human c-Src kinase domain phosphorylation of a preferred peptide substrate, PKCδ-Y313 (SSEPVGI**Y**QGFEKKT) and a non-preferred peptide, PDGFRα-Y762 (SDIQRSL**Y**DRPASAK), in which the Tyr 768 in the wildtype PDGFRα sequence was mutated to alanine in the peptide so as to present a single substrate tyrosine residue, were used as controls in these kinase assays [17]. The tyrosine phosphorylation kinetics for RTKC8 with these two previously mentioned peptides and three peptides from the RTKC8 C-terminal tail (purchased from Elim Biopharmaceuticals, Inc.), corresponding to the residues surrounding Tyr 1828 (EPETDEV**Y**GNEGNVS), Tyr 1897 (GTVGEHE**Y**FDCADQH), and Tyr 1990 (ASDNELL**Y**DMGRAEA) were measured. The reactions were initiated by the addition of either c-Src at 1 µM or RTKC8 kinase domain at 5 µM, to 250 mM peptide. The reaction progress was monitored by measuring absorbance at 340 nm every 30 s at 25°C on a SpectraMax plate reader.

### Crystallization and data collection

The freshly purified RTKC8 kinase domain (10-15 mg/mL, in Buffer C) was mixed with a final concentration of 1 mM staurosporine (10 mM stock, dissolved in 100% DMSO) and incubated on rotator for 2 hours at room temperature. The crystallization trays (96 well) were set up using sitting-drop vapor diffusion with a Mosquito crystallization robot (TTP Labtech) using commercial sparse matrix screens. 0.1 µL of the protein was mixed with 0.1 µL of the reservoir solution. The drops were equilibrated against 50 µL of reservoir volume and the trays were incubated at 20°C. A condition from the PACT screen (Qiagen) yielded crystals suitable for diffraction (0.1 M succinic acid, sodium dihydrogen phosphate and glycine (SPG) buffer pH 6.0, 25% (w/v) polyethylene glycol (PEG) 1500 [18]. The crystals were cryoprotected with a solution of the well condition with 20% glycerol (v/v) and X-ray diffraction data were collected at Lawrence Berkeley National Laboratory Advanced Light Source, Beamline 8.2.1 with 0.9998 Å wavelength X-rays.

### Structure Determination

The reflections were indexed and integrated by XDS [19,20]. The data were scaled and merged with Pointless [21] and Aimless [22], in the CCP4 suite [23]. The structure was solved by molecular replacement with Phaser-MR in the Phenix suite [24] using the structure of tyrosine kinase AS - a hypothetical protein with a sequence corresponding to a common ancestor of Src and Abl with ∼ 43% identity to the RTKC8 kinase domain (PDB 4UEU [25]) as a search model. The structure was rebuilt using Phenix-AutoBuild and refined with multiple cycles of refinement with phenix.refine [26] and model building with Coot [27]. The calculated difference Fourier map (Fo-Fc) revealed clear electron density for two molecules of bound staurosporine per kinase domain. The coordinates and structure factors of the final structure have been deposited in the Protein Data Bank (PDB). Software used in this project was curated by SBGrid [28].

## Results

### RTKC8 kinase domain is phosphorylated

Western blot analysis was performed to check the phosphorylation status of the purified RTKC8 kinase. The immunoblot was probed with an anti-phosphotyrosine antibody and showed that the purified RTKC8 kinase had at least one phosphorylated tyrosine residue which could be dephosphorylated by the tyrosine-protein phosphatase YopH (Fig 2A). The band intensity was decreased to ∼30% of the initial level after five minutes and to less than ∼5% after an hour.

**Fig 2.**
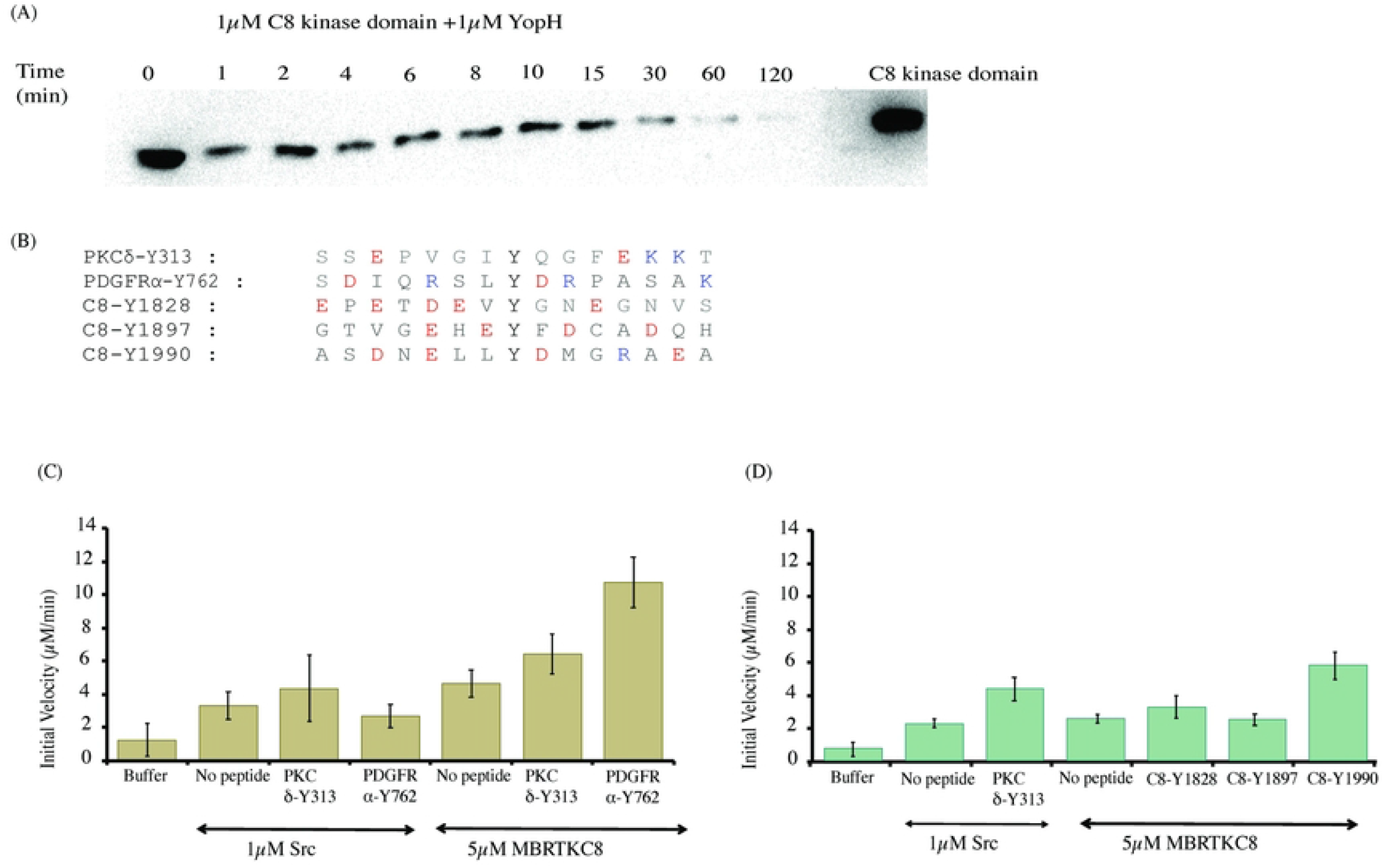
Phosphorylated RTKC8 kinase domain is active and phosphorylates tyrosine residues in tail segment peptides. (A) The RTKC8 kinase domain was dephosphorylated by YopH phosphatase and samples were taken at different time intervals and probed with an anti-phosphotyrosine antibody. The control showed that the band intensity on the western blot for the kinase domain without YopH is unchanged over 2 hours at room temperature while the band intensity decreased rapidly upon the addition of the tyrosine phosphatase YopH. (B) The sequence of tyrosine phosphosite peptides (PKCδ-Y313, PDGFRα-Y762, RTKC8-Y1828, RTKC8-Y1897, RTKC8-Y1990) used as substrates for the kinase (C) Phosphorylation activity of RTKC8 kinase domain on the tyrosine containing peptides PKCδ-Y313 and PDGFRα-Y762 (D) Phosphorylation activity of RTKC8 kinase on tyrosine residue containing peptides from the tail segment (RTKC8-Y1828, RTKC8-Y1897, RTKC8-Y1990).

### Substrate specificity of the RTKC8 kinase

The phosphorylation rates of the peptides human c-Src preferred peptide (PKCδ-Y313) and non-preferred peptide (PDGFRα-Y762) [17] by c-Src (1 µM) and RTKC8 kinase domain (5 µM) were assayed (Figs 2B and C). RTKC8 kinase appears to have a different substrate specificity than c-Src, showing more rapid phosphorylation of PDGFRα-Y762 than PKCδ-Y313. Kinase assays were also performed with three RTKC8 tail peptides comprised of the sequences surrounding Tyr 1828, Tyr Y1897, or Tyr Y1990. These experiments showed that the different tail peptides were phosphorylated at different rates. The RTKC8 peptide containing Tyr 1990 had the highest rate of phosphorylation by RTKC8 kinase (Figs 2B and D). Both the c-Src non-preferred peptide (PKCδ-Y313) and the tail peptide (RTKC8-Y1990) have Leu before the Tyr and Asp after the phosphosite tyrosine residue. This result suggests that the preferred substrates of the RTKC8 kinase contain a hydrophobic residue at the position immediately before the phosphosite and an acidic residue immediately after it.

### Structure determination

Crystallization trials of the RTKC8 kinase domain with staurospaurine yielded triangular shaped crystals, 70-100 µm in diameter (Fig 3A). The crystals were of the orthorhombic space group P2_1_2_1_2 with cell dimensions a = 81.1, b = 55.5, c = 60.6 (Å). The crystal structure of the kinase domain was determined to 1.95 Å resolution and the model was built with a final Rwork/Rfree (%) of 20.4/25.7 (Table 1). The crystallographic model includes residues 1532 to 1804, two staurosporine molecules, ninety water molecules and one phosphate group. The first three residues (1529-1531), the last seven residues (1805-1811) and a region of the activation loop (1694-1698) in the expressed protein construct were not modeled as there was no interpretable electron density for these residues. The tyrosine residues which would be expected to be phosphorylated in the activation loop (Tyr 1697 and Tyr 1698) were in the disordered activation loop region. The staurosporine inhibitor bound to the kinase domain at two sites, one in the ATP binding site and other on the C-lobe in the peptide-substrate binding site.

**Fig 3.**
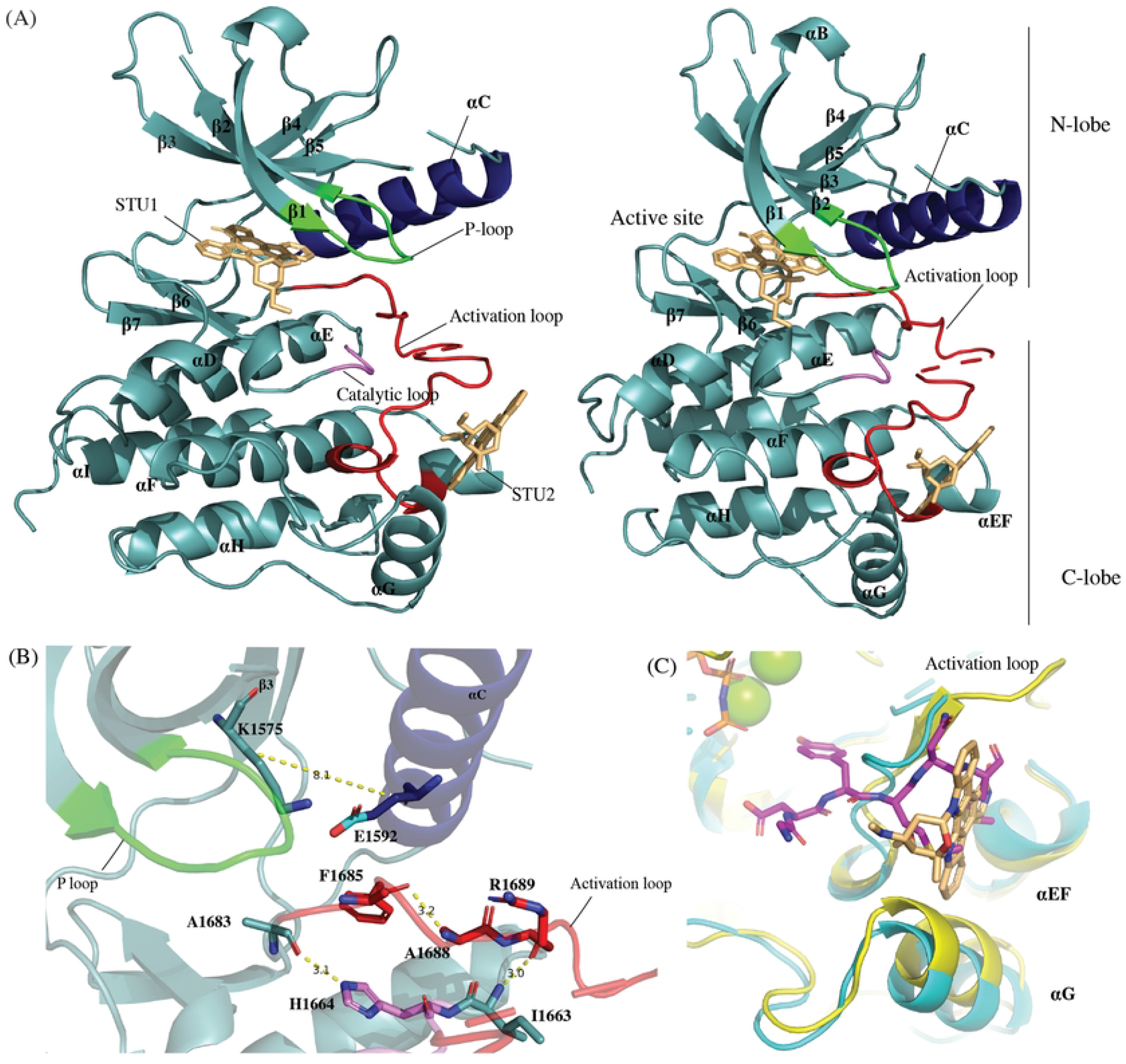
The RTKC8 kinase domain crystallized in an active conformation with staurosporine. (A) The RTKC8 structure reveals a classic, two lobed kinase domains (cyan) with an αC helix (blue), P-loop (green), activation loop (red), catalytic loop (violet) with two staurosporines bound, one at the active site and the other at the peptide-substrate binding site (light orange). (**B**) The active αC-helix-in conformation is demonstrated by the presence of a salt bridge between the conserved αC-helix Glu1592 and Lys1575 in the β3 strand (distance between the two Cβs is <10Å), the presence of a hydrogen bond between the backbone of the amino acid immediately prior to the DFG and the His in the HRD motif and an extended activation loop with a hydrogen bond between N-terminus of Arg1689 of the DFGxxR motif and the backbone carbonyl of Ile1663 which is immediately before the HRD motif. A hydrogen bond in the extended activation loop between the DFG motif Phe1685 and DFGxA, Ala1688 presents another feature of an active conformation of a kinase. (C) The second staurospaurine binds at the peptide-substrate binding site. When the structure of the human insulin kinase domain (yellow) in complex with a peptide substrate (1IR3) is overlayed on the RTKC8 kinase domain (cyan), the second staurospaurine (light orange) can be seen overlapping the position of the C-terminal segment of the substrate peptide (magenta) bound between the activation loop, the αEF helix and the αG helix.

**Table 1.** Structure determination and refinement of RTKC8 kinase domain bound to staurosporine.

|  |  |
| --- | --- |
| Data Collection |  |
| Wavelength (Å) | 0.9998 |
| Space group | P2<sub>1</sub>2<sub>1</sub>2 |
| Cell dimensions (Å) | a = 81.1, b = 55.5, c = 60.6 |
| Resolution (Å) | 48.55-1.95 |
| Rmerge | 0.23(3.648) |
| Rmeas | 0.235(3.758) |
| Rpim | 0.046(0.883) |
| Total Observations | 539294(23983) |
| Unique Observations | 20593(1414) |
| CC1/2 | 0.998(0.455) |

|  |  |
| --- | --- |
| Completeness (%) | 99.9(99.3) |
| Multiplicity | 26.2(17.0) |
| $I/\sigma(I)$ | 12.4(0.9) |
| Refinement Statistics |  |
| Rfree test set | 1039 |
| Resolution | 48.55-1.95 |
| R-factor (%) | 20.39 |
| Rfree (%) | 25.72 |
| No. of Atoms |  |
| Protein | 2167 |
| Ligand | 75 |
| STU | 70 |
| PO<sub>4</sub> | 5 |
| Water | 90 |
| Average B-factor |  |
| Proteins | 36.03 |

|  |  |
| --- | --- |
| Ligand | 38.54 |
| Ramachandran Statistics |  |
| Favoured (%) | 97.35 |
| Disallowed (%) | 0.00 |
| Root mean square deviation from ideality |  |
| Bonds (Å) | 0.007 |
| Angle (°) | 0.855 |
| MolProbity clash score | 2.08 |

### The choanoflagellate RTKC8 kinase domain crystallized in an active conformation

The choanoflagellate kinase domain exhibits the conserved structural features of a typical kinase domain [29,30] with two lobes, the N-lobe and C-lobe. The activation loop is a stretch of 29 amino acids in the C-lobe, beginning with the conserved Asp-Phe-Gly (DFG) motif and ending at Ala-Pro-Glu (APE). The activation loop of the RTKC8 kinase domain contains two consecutive tyrosine residues for which there is no observable electron density.

The RTKC8 kinase crystallized in an active conformation with a RMSD of 1.0 Å over 201 Cα atoms to the structure of Lck in an active conformation [31]. The conserved Glu 1592 of the αC-helix is rotated towards the active site to form a salt bridge with Lys 1575 of the β3-strand, oriented as “αC-helix-in”. The distance between αC-helix-Glu-Cβ and β3-Lys-Cβ atoms is 8.1 Å (C-helix-in and C-helix-out structures are defined with the distance of ≤10 Å or >10 Å, respectively) [32]. The kinase is crystallized in “DFGin” conformation where the DFG-Asp is pointed towards the active site and the DFG-Phe interacts with the αC-helix. The Asp backbone and its sidechain is oriented in such a manner that the carbonyl group of the Ala prior to the DFG motif forms a hydrogen bond with the nitrogen of the histidine sidechain in HRD motif [32]. The distance between the sidechain nitrogen of His 1664 and the carbonyl group of the Ala 1683 is 3.1 Å, consistent with formation of a hydrogen bond between them. The activation loop is extended away from the C-lobe of kinase domain. In this extended conformation, the cleft is accessible to ATP and peptide substrates [33][34]. The orientation of the activation loop leads to the formation of a hydrogen bond between the backbone N atom of the sixth residue in the loop (DFGxx**X)** and the backbone carbonyl of the residue prior to the HRD motif in the catalytic loop. The amide nitrogen of Arg 1689 (DFGxx**R**) and the carbonyl of Ile 1663 form a hydrogen bond of 3 Å. Another major structural feature of this conformation is the presence of type I β-turn at the Phe in the DFG motif, which forms a hydrogen bond between the carboxyl group of the Phe and the NH of the Ala two residues after the DFG motif [32]. The hydrogen bond length between Phe 1685 and Ala 1688 is 3.2 Å. This interaction plays an important role in orientating the activation loop to an extended conformation. All these interactions and structural features present the signature of the active conformation of the kinase domain (Fig 3C).

### Staurosporine interactions with the RTKC8 kinase domain

Two molecules of staurosporine are bound to the RTKC8 kinase. One is in the ATP-binding site as seen in several other structures of kinase:staurospaurine structures (eg. CDK2 (1AQ1) and ZAP-70 (1U59) (Fig 3A), and the other is bound to the activation loop and at the C-lobe (Figs 2C and 3B). The active site inhibitor is positioned in the large groove surrounded by β strands in the N-lobe, the glycine rich P-loop, the hinge region, the N-terminus of the activation loop and the small helix between αE and β6 of the C-lobe (Fig 4A). There are both hydrophobic and hydrophilic interactions between the inhibitor and residues inside the cleft. There are three hydrogen bonds, with Glu 1622, Met 1624 and Arg 1670 and five hydrophobic interactions, with Leu 1549, Val 1557 and Ala 1573 from N-lobe, and Gly 1627 and Leu 1673 from C-lobe. The hydrogens from the secondary amine group and the keto oxygen of the lactam ring of the inhibitor make a pair of hydrogen bonds with carbonyl oxygen on the backbone of Glu1622 and the primary amine on the backbone of Met1624 in the hinge region, respectively. The third hydrogen bond is observed between the methyloamino nitrogen of staurosporine and the carbonyl oxygen on the backbone of Arg1670, in the ribose-binding pocket of the kinase domain (Fig 4B).

**Fig 4.**
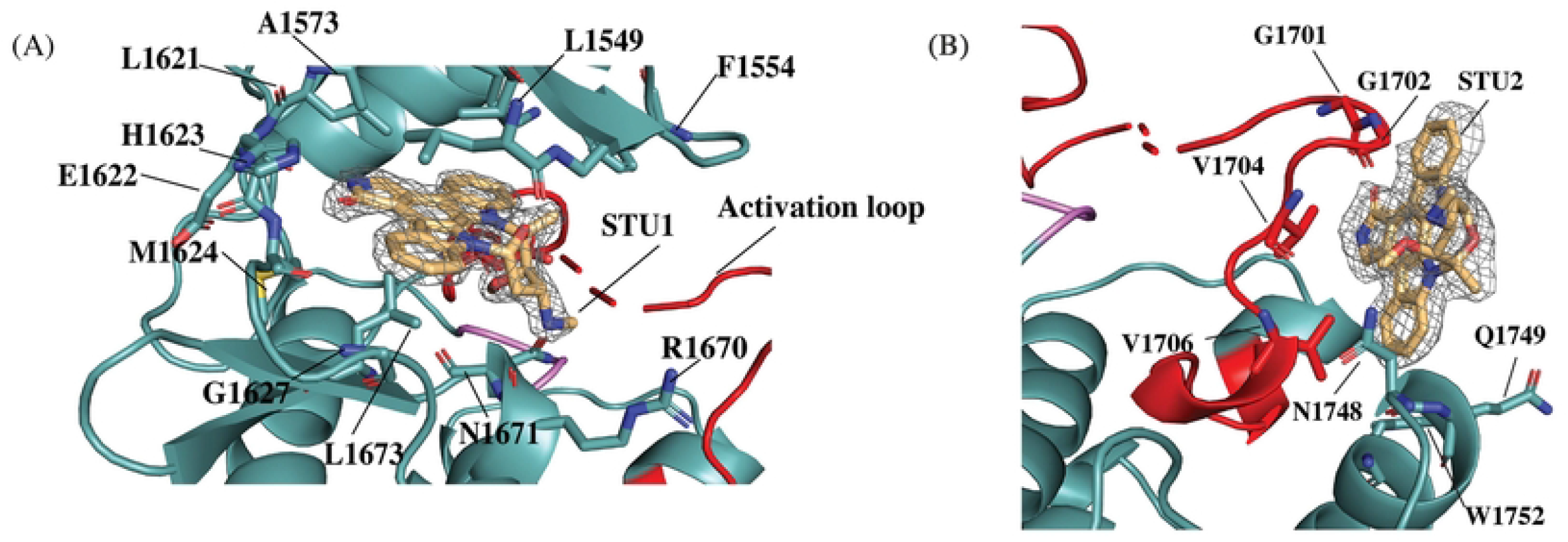
Two molecules of staurosporine (STU1 and STU2) are bound to the RTKC8 kinase. (A) One molecule of staurosporine (STU1) is bound at the active site, surrounded by the N-lobe β strands, the activation loop and the small helix between αE and β6 of the C-lobe (electron density for the Fo-Fc map with no compound modeled contoured at 2.0σ in grey). (B) The other staurosporine (STU2) is bound to the C-lobe, next to the activation loop, the αEF helix and the αG helix (electron density for the Fo-Fc map with no compound modeled contoured at 2.0σ in grey).

Another molecule of staurosporine was found bound to the C-lobe of the kinase domain in a hydrophobic groove between the activation loop, the αEF helix and the αG helix (Fig 4C). This staurosporine is bound at the peptide-substrate binding site as revealed in the structure of the insulin receptor tyrosine kinase domain in complex with a peptide substrate (1IR3) [35] (Fig 3C). This site has been identified as a druggable site, [36] and is the site of a trans interaction in Aurora kinases, where the activation loop from one kinase binds to and presents as a substrate for phosphorylation by the other aurora kinase, so is designated the AAS site. This potential binding site has been identified in Src, Zap-70, and other tyrosine kinases using solvent mapping algorithms [36], and it is intriguing to see the inhibitor bound to a tyrosine kinas AAS site. The ligand interacts with hydrophobic residues (Gly 1701, Gly 1702, Val 1704, Val 1706) in the activation loop and Trp 1752, Asn 1748 and Gln 1749 in the αG-helix (Fig 4D).

### Structural comparison of the RTKC8 kinase domain with the Ephrin kinase domain

A Dali search [37] of the RTKC8 kinase model (268 residues) from *Choanoflagellate* against the PDB database shows the highest structural similarity with the kinase domain of the Ephrin type A receptor 5 (2R2P, 295 residues) with the z-score of 36.2 and 38% sequence identity. The two structures were aligned over 214 atoms with a root mean square deviation of 0.84 Å. (Fig 5). The Ephrin receptor juxtamembrane region modulates the kinase activity [38]. Two tyrosine residues in the juxtamembrane segment, when phosphorylated, release inhibition, and provide docking sites for SH2 domain-containing proteins to transmit the downstream signal [39]. However, the juxtamembrane domain of RTKC8 kinase does not contain tyrosine residues. In addition, while the kinase domains are structurally similar, the extracellular receptor on the Ephrin receptor bears no resemblance to that of the RTKC8 receptor. The structure of RTKC8 kinase demonstrates that the kinase domain remained highly conserved throughout the evolution of receptor tyrosine kinases from choanoflagellates to human but with divergent fused extracellular domains.

**Fig 5.**
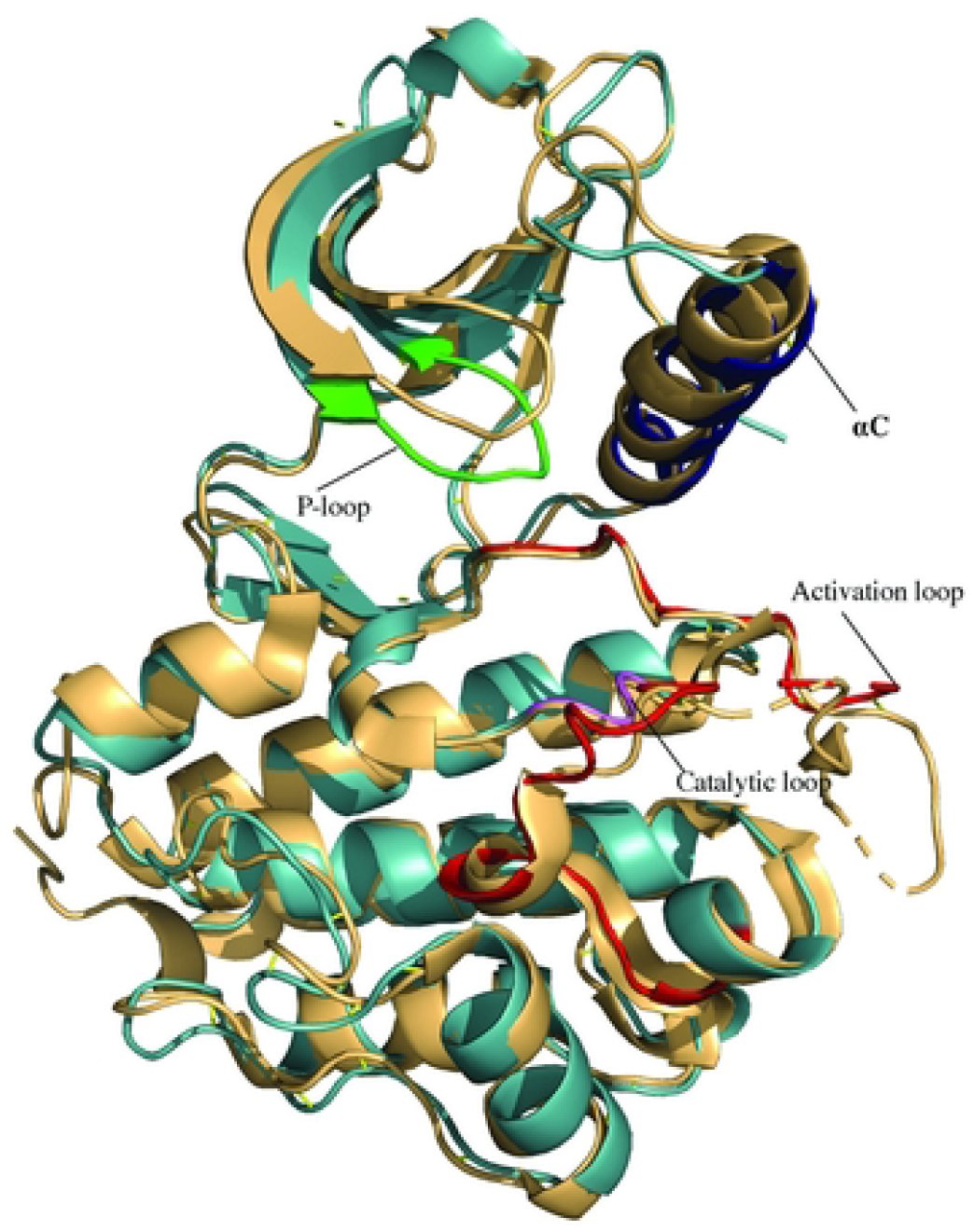
Structure alignment of RTKC8 kinase and EphA5 kinase. Dali search identified the kinase domain of EphA5 (2R2P) as the structure with the highest similarity in the PDB. The RTKC8 kinase (cyan) and EphA5 kinase (yellow) structures were aligned with a rmsd 0.836 Å.

## Concluding Remarks

Choanoflagellates provide an excellent opportunity to study the evolution of receptor tyrosine kinases as they are the only group shown to have receptor tyrosine kinases outside of metazoans [9]. We report the expression, purification, and initial characterization of the RTKC8 kinase domain from a choanoflagellate receptor tyrosine kinase. We also present the crystal structure of the kinase domain in complex with staurospaurine. The purified RTKC8 kinase is catalytically active and can phosphorylate tyrosine residue containing peptides generated from its C-terminal tail sequence which further suggests that the tail tyrosine residues act as substrates. The secondary binding site for staurospaurine in the substrate binding site shows the potential for this site in developing tyrosine kinase inhibitors.

## Acknowledgements

We thank Xiaoxian Cao of the Kuriyan Lab for initial assistance with protein expression in insect cell culture. We thank Nicole King for providing the Monosiga cDNA library. We thank the Berkeley Center for Structural Biology beamline staff at the Advanced Light Source, Lawrence Berkeley National Laboratory. The Berkeley Center for Structural Biology is supported in part by the Howard Hughes Medical Institute. The Advanced Light Source is a Department of Energy Office of Science User Facility under Contract No. DE-AC02-05CH11231. We are thankful to Ruchika Bajaj and Neel H. Shah for providing comments and suggestions to improve the manuscript.

